# Deep assessment of human disease-associated ribosomal RNA modifications using Nanopore direct RNA sequencing

**DOI:** 10.1101/2021.11.10.467884

**Authors:** Isabel S. Naarmann-de Vries, Christiane Zorbas, Amina Lemsara, Maja Bencun, Sarah Schudy, Benjamin Meder, Jessica Eschenbach, Denis L.J. Lafontaine, Christoph Dieterich

## Abstract

The catalytically active component of ribosomes, rRNA, is long studied and heavily modified. However, little is known about functional and pathological consequences of changes in human rRNA modification status. Direct RNA sequencing on the Nanopore platform enables the direct assessment of rRNA modifications. We established a targeted Nanopore direct rRNA sequencing approach and applied it to CRISPR-Cas9 engineered HCT116 cells, lacking specific enzymatic activities required to establish defined rRNA base modifications. We analyzed these sequencing data along with wild type samples and *in vitro* transcribed reference sequences to specifically detect changes in modification status. We show for the first time that direct RNA-sequencing is feasible on smaller, i.e. Flongle, flow cells. Our targeted approach reduces RNA input requirements, making it accessible to the analysis of limited samples such as patient derived material. The analysis of rRNA modifications during cardiomyocyte differentiation of human induced pluripotent stem cells, and of heart biopsies from cardiomyopathy patients revealed altered modifications of specific sites, among them pseudouridines, 2’-O-methylation of riboses and acetylation of cytidines. Targeted direct rRNA-seq analysis with JACUSA2 opens up the possibility to analyze dynamic changes in rRNA modifications in a wide range of biological and clinical samples.

## Introduction

Ribosomes as “protein synthesis machineries” are essential components of every cell and largely conserved throughout all three kingdoms of life. They are composed of a large and a small subunit, in eukaryotes designated as 60S and 40S subunit, respectively. In mammals, the ribosome is composed of four ribosomal RNAs (5S, 5.8S, 18S, 28S) and 80 ribosomal proteins (*1*). The biogenesis of ribosomes is a highly complex process that requires several hundreds non-ribosomal factors (*2–5*). These non-ribosomal factors can be grouped into protein (complexes) and snoRNPs.

One important step of faithful ribosome biogenesis is the modification of ribosomal RNAs (rRNAs) (*6, 7*) with some of these modifications being conserved from bacteria to eukaryotes. Most frequently found modifications are the 2’-O-methylation of ribose and isomerization of uridine to pseudouridine (*8*). In a comprehensive study on 18S rRNA 43 pseudouridines and 42 ribose methylations were described, whereas the 28S rRNA was shown to contain 61 pseudouridines and 68 ribose methylations (*8*). Besides these frequent modifications, rRNAs are also modified at individual bases by methylation, acetylation and aminocarboxypropylation (*8*). For most of the ribosomal modifications the catalyzing enzymes have been identified, however their biological role is often only partially understood. 2’-O-methylations and pseudouridylations that are catalyzed by snoRNA guided enzyme complexes (*9*) are thought to stabilize secondary and tertiary structures of modified rRNAs (*10–12*). Furthermore, they are required to maintain efficient and proper translation (*12–14*).

The importance of correct ribosome assembly is evidenced by a class of diseases that are designated as “ribosomopathies” and frequently caused by mutations in different ribosomal proteins or factors required for ribosome biogenesis (*15*). A prominent example is Diamond Blackfan anemia that is caused by mutations in different ribosomal proteins (*16*). Although all ribosomopathies affect the universal protein synthesis machinery, they give rise to different mutation-specific disease entities (*15*). Mutations in the pseudouridine synthase enzyme DKC1 (dyskerin) cause X-linked dyskeratosis congenita (*17*), which is characterized by failure of proliferating tissues including skin, mucosa and bone marrow and increased cancer susceptibility, indicating that also aberrant RNA modifications can contribute to disease. The detected decrease in pseudouridines is accompanied by an impairment in the translation of specific cellular mRNAs bearing internal ribosome entry sites in their 5’UTR (*12*). Furthermore, translational fidelity is strongly reduced by DKC1 depletion (*18, 19*).

Recently, a *trans* locus was identified that causes misregulation of SNORA48 (predicted to catalyze a pseudouridylation on the 28S rRNA (*20*)) and ribosomopathy in hypertrophic hearts (*21*). Furthermore, BUD23/WBSCR22 (catalyzing 18S m^7^G_1639_ (*22–24*)) was shown to be required for efficient translation of mitochondrial transcripts and its deletion causes severe cardiomyopathy (*25*). These recent findings indicate that changes in rRNA modification might be implicated in cardiovascular diseases as well, however until now differences in modification levels have not been demonstrated directly.

In the past, the analysis of mutations in ribosomal proteins and ribosome biogenesis processing factors was possible. However, it was not possible to systematically elucidate mutations in rRNA (due to the high complexity of genomic rRNA loci (*26, 27*)) or directly assess dynamic changes in distinct rRNA modifications in patient samples for which the amount of material is limited. Mass spectrometry- and HPLC-based methods usually require a high amount of input material (20 μg total RNA) to allow the accurate detection of ribosomal modifications (*8*) and other methods are restricted to specific modification types. Consequently, the relevance of mutations or aberrant modifications in ribosomal RNAs are not well understood. With the advent of long read sequencing techniques, especially the direct RNA sequencing method introduced by Oxford Nanopore Technologies (ONT), it is for the first time possible to directly sequence full length RNA molecules (*28–30*). Using Nanopore direct RNA-seq, also RNA modifications can now be analyzed in a direct and specific manner, so far mainly assessed for m^6^A (*28,29,31-34*). Nanopore direct RNA sequencing (direct RNA-seq) was previously used to analyze rRNA sequences derived from *E.coli* (*32,33*), yeast (*33,35*) and human cells (*33*). Several computational tools are available to identify RNA modifications in Nanopore direct RNA-seq data (*33,35-38*) with different underlying concepts. Notably, they can be based either on basecalling errors or changes in the raw signal. Furthermore, some algorithms depend on training data, whereas others are designed for comparative analysis. We have recently introduced JACUSA2 for the analysis of RNA modifications (*34*), which uses basecalling errors (Mismatch, Deletion, Insertion) in pairwise comparisons (call-2) and handles replicate samples.

Here, we established targeted direct RNA-seq of human 18S and 28S rRNA (Nanopore direct rRNA-seq) and employed a set of CRISPR-Cas9 engineered human HCT116 cells lacking specific rRNA modifications on the 18S rRNA to detect modification signatures in Nanopore direct rRNA-seq data based on the comparative analysis with wild type (WT) and *in vitro* transcribed (IVT) rRNA. Our analysis of these genetic model systems was focused on specific 18S rRNA base methylations, namely METTL5 catalyzed m^6^A modification of A_1832_ (*39*) and DIMT1L installed dual dimethylation of A_1850_A_1851_ (*24, 40, 41*). Furthermore, we analyzed the methylation of m^7^G_1639_, introduced by BUD23/WBSCR22 (*22–24*). Employing JACUSA2 call-2 (pairwise comparison) we show that all analyzed rRNA modifications are detectable in Nanopore direct rRNA-seq data. To expand the repertoire of biological samples that are accessible to Nanopore direct rRNA-seq, the targeted rRNA sequencing was transferred to recently introduced Flongle flow cells. We demonstrate for the first time that direct RNA-seq is feasible on these small devices equipped with 126 pores and show that downscaled Nanopore direct rRNA-seq allows comparable detection of RNA modifications. Following up on these promising results, we sampled different read numbers from our sequencing data, and show that detection of the analyzed base modifications is possible with as little as 300-500 reads. Furthermore, we provide evidence that the level of modification can be estimated based on a calibration curve.

To illustrate our direct rRNA-seq - JACUSA2 framework accomplishs characterization of dynamic rRNA modifications, we analyzed the rRNA modification status in the course of cardiac differentiation of human induced pluripotent stem cells (hiPSCs). Importantly, this setup can be applied to the study of patient derived samples as well. Notably, our analysis of human heart biopsies revealed ac^4^C_1842_ as a candidate site for differential modification in cardiovascular diseases, such as dilated cardiomyopathy and hypertrophic cardiomyopathy, making it a potential biomarker.

## Results

### Targeted direct rRNA-seq in total human RNA

The direct analysis of rRNA sequence variants and rRNA modifications in limited amounts of input material is just now becoming possible, mainly due to the advent of the direct RNA-seq platform developed by Oxford Nanopore Technologies (ONT). Our aim was to establish a protocol with minimal pre-processing and requirement of input material to enable the analysis of low-input samples and patient-derived material. To prevent the laborious purification of individual rRNAs, which is often complicated by material loss, or additional experimental steps, such as *in vitro* polyadenylation, we established custom adapters for the sequencing of human 18S and 28S rRNA. Employing *in vitro* transcribed (IVT) 18S rRNA, we compared the performance of two custom adapters with different lengths, 18S v1 (10 nts specific sequence) and 18S v2 (20 nts specific sequence) (in blue, Fig. 1A) with the standard oligo(dT) adapter (RTA) after polyadenylation of the IVT. Direct RNA-seq libraries were prepared with the three different adapters and analyzed on MinION (R9.4.1) flow cells. Median read length and quality of reads were slightly higher in sequencing runs employing the custom adapters (Fig. 1B). Furthermore, we noticed a higher coverage at the 3’end of the IVT in libraries prepared with the custom adapters (Fig. 1C). As ten nucleotides were sufficient to target the 18S rRNA, we decided to use the 18S v1 adapter for subsequent experiments and designed an analogous 28S rRNA adapter (see Material and Methods). These adapters were then used for direct RNA-seq of rRNA from human HCT116 cells and compared to the sequencing of full length 18S IVT or a 1452 nt long 3’ IVT fragment of the 28S rRNA (Fig. 1D,E). Both custom adapters specifically target their respective rRNA, however sequencing of the 28S rRNA was about ten-fold less efficient in terms of obtained reads and especially full length reads (Fig. 1E). This is likely explained by the presence of long homo-polymeric, very GC-rich stretches on the 28S rRNA, which are known to cause handling problems for the Nanopore (*42, 43*). The cellular rRNA displayed a higher mismatch rate than the IVT (colored lines, Fig. 1D,E), which is indicative for the presence of rRNA modifications that can by detected via basecalling errors with JACUSA2 (*34*). We compared HCT116 rRNA with the respective IVTs using JACUSA2 call-2 in pairwise comparisons and considered the Scores for Mismatch, Deletion and Insertion. Indeed, most previously described modifications sites (*8*) are represented by high JACUSA2 scores (Fig. 1F). In conclusion, Nanopore direct rRNA-seq is suitable for the detection of modifications on human rRNAs.

**Figure 1:**
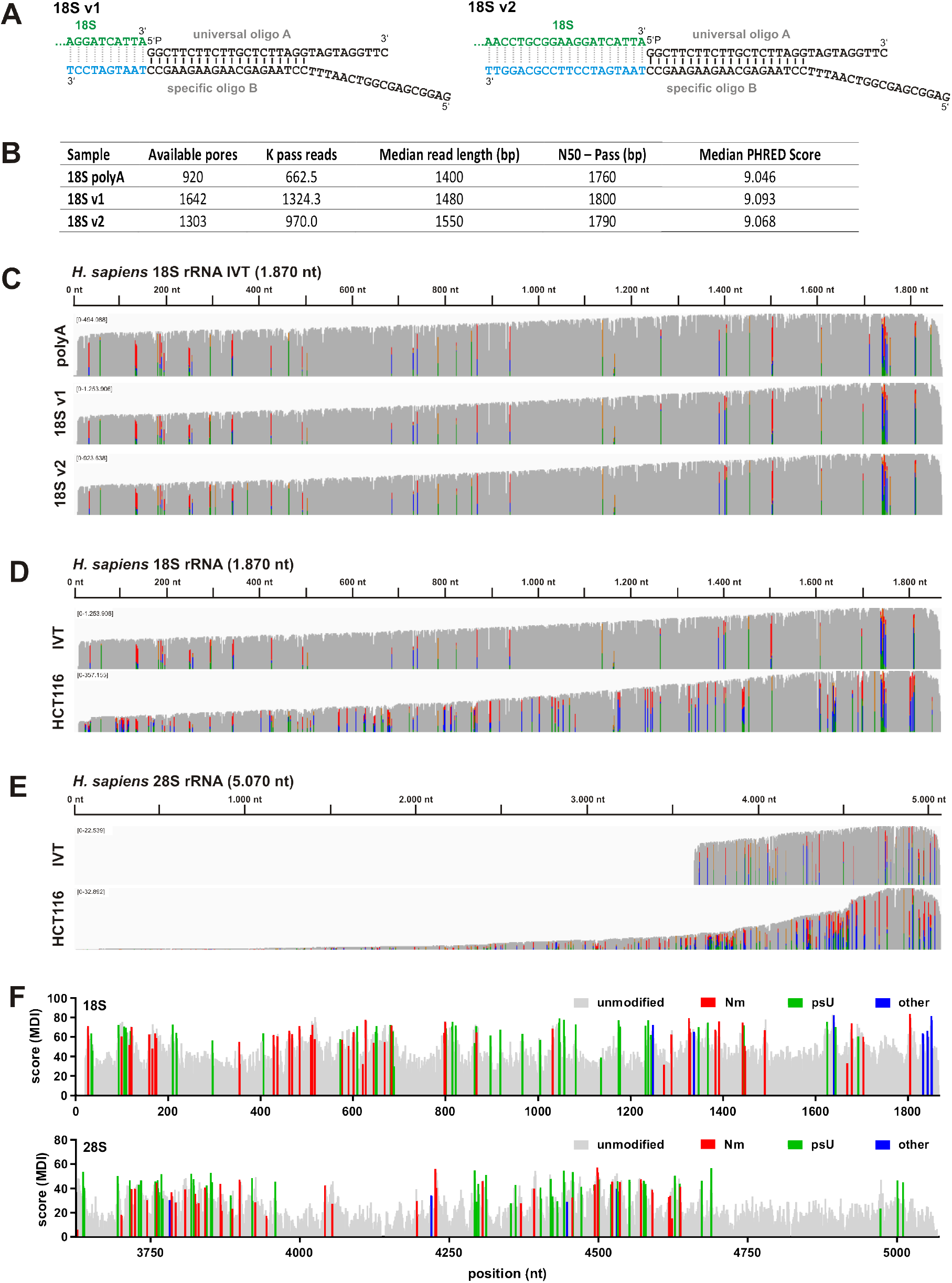
Setup of targeted direct rRNA sequencing. A) Schematic representation of tested custom adapters, 18S v1 and 18S v2. The 3’end of the 18S rRNA is depicted in green. The custom adapters consist of an universal oligo A annealed to a sequence specific oligo B. The sequence specific part of oligo B (light blue) has a length of either 10 nts (18S v1) or 20 nts (18S v2). B) Read statistics of Nanopore direct RNA-seq of 18S IVT with the standard polyA adapter (after polyadenylation of the IVT), 18S v1 and 18S v2 on MinION R9.4.1 flow cells. C) Coverage of 18S IVT from IGV snapshots of the sequencing runs listed in B). D) Coverage of Nanopore direct rRNA-seq of 18S IVT and 18S rRNA from HCT116 WT cells sequenced on MinION R9.4.1 flow cells. E) Coverage of Nanopore direct rRNA-seq of 28S 3’ fragment IVT and 28S rRNA from HCT116 WT cells sequenced on MinION R9.4.1 flow cells. C-E) Allele frequency threshold = 0.2. Mismatches are indicated by color. F) JACUSA2 call-2 analysis of HCT116 WT vs. IVT for 18S (top) and 28S 3’fragment (bottom) considering the Mismatch, Deletion and Insertion scores for each site (MDI). Known modification sites are labeled according to the modification type as indicated.

### Nanopore direct rRNA-seq enables detection of site specific RNA modifications

Modification sites on rRNA, and also human rRNA, are well characterized by a range of techniques, including classical RNA biochemistry, HPLC, mass spectrometry- and short read deep sequencing-based methods (see Introduction). For the large majority of these modifications, responsible enzymes and snoRNAs that catalyze site-specific modifications have been identified and characterized. We made use of this knowledge to analyze specific rRNA modifications in human cells lacking individual modifications by targeted Nanopore direct RNA-seq with a limited amount of input material.

For this, we used human HCT116 derived cell lines that were genetically engineered by CRISPR-Cas9 to harbor either a knock out (KO) or a catalytic-dead variant (MUT) of selected methyltransferases (see Material and Methods). The cell lines and affected modifications are listed in Table 1. We compared in each set IVT, WT and KO/MUT, which were sequenced on standard MinION (R9.4.1) flow cells and basecalled with the MinKNOW-embedded Guppy basecaller. Reads that mapped to the human 18S rRNA were analyzed for the presence of rRNA modifications employing the recently introduced JACUSA2 framework (*34*) in pairwise comparisons (call-2). We tested different Feature sets from JACUSA2 to account for the clustering of rRNA modifications, as well as for the inherent characteristic of Nanopore sequencing to read sequences in 5-mers. Initially, we compared four different Feature sets: a) Mismatch score of the analyzed site (M), b) Mismatch, Insertion and Deletion scores of the analyzed site (MDI), c) Mismatch score of the 5-mer context (modified site in position 3), Insertion and Deletion score of the analyzed site (M_Con_DI), d) Mismatch, Deletion and Insertion score in the 5-mer context ((MDI)_Con_).

**Table 1:** Genetically engineered HCT116 cell lines analyzed by direct rRNA-seq

| cell line | missing modification | reference modification enzyme |
| --- | --- | --- |
| HCT116 wt | - |  |
| HCT116 WBSCR22<sup>D82K/D82K</sup> | 18S m<sup>7</sup>G<sub>1639</sub> | (22–24) |
| HCT116 METTL5<sup>-/-</sup> | 18S m<sup>6</sup>A<sub>1832</sub> | (39, 56) |
| HCT116 DIMT1L<sup>Y131G/Y131G</sup> | 18S m<sup>6</sup><sub>2</sub>A<sub>1850</sub> m<sup>6</sup><sub>2</sub>A<sub>1851</sub> | (24, 40, 41) |

The comparison of the different Feature sets revealed that the Mismatch score alone is not sufficient to discriminate target from non-target sites (Figure S1). Consideration of Deletion and Insertion Scores (MDI) improved detection (Figure S1) and identified all target sites as significant outliers in the MinION sequencing data. The Feature set composed of the Mismatch Score for the 5-mer context (target site plus nts −2 to +2), Deletion and Insertion Score for the modification site (M_Con_DI), turned out to be most robust for the detection of the strong and sitespecific differences in rRNA modification represented by the genetic models and is displayed in Fig. 2D-F. All other Feature sets tested can be found in Figure S1.

**Figure 2:**
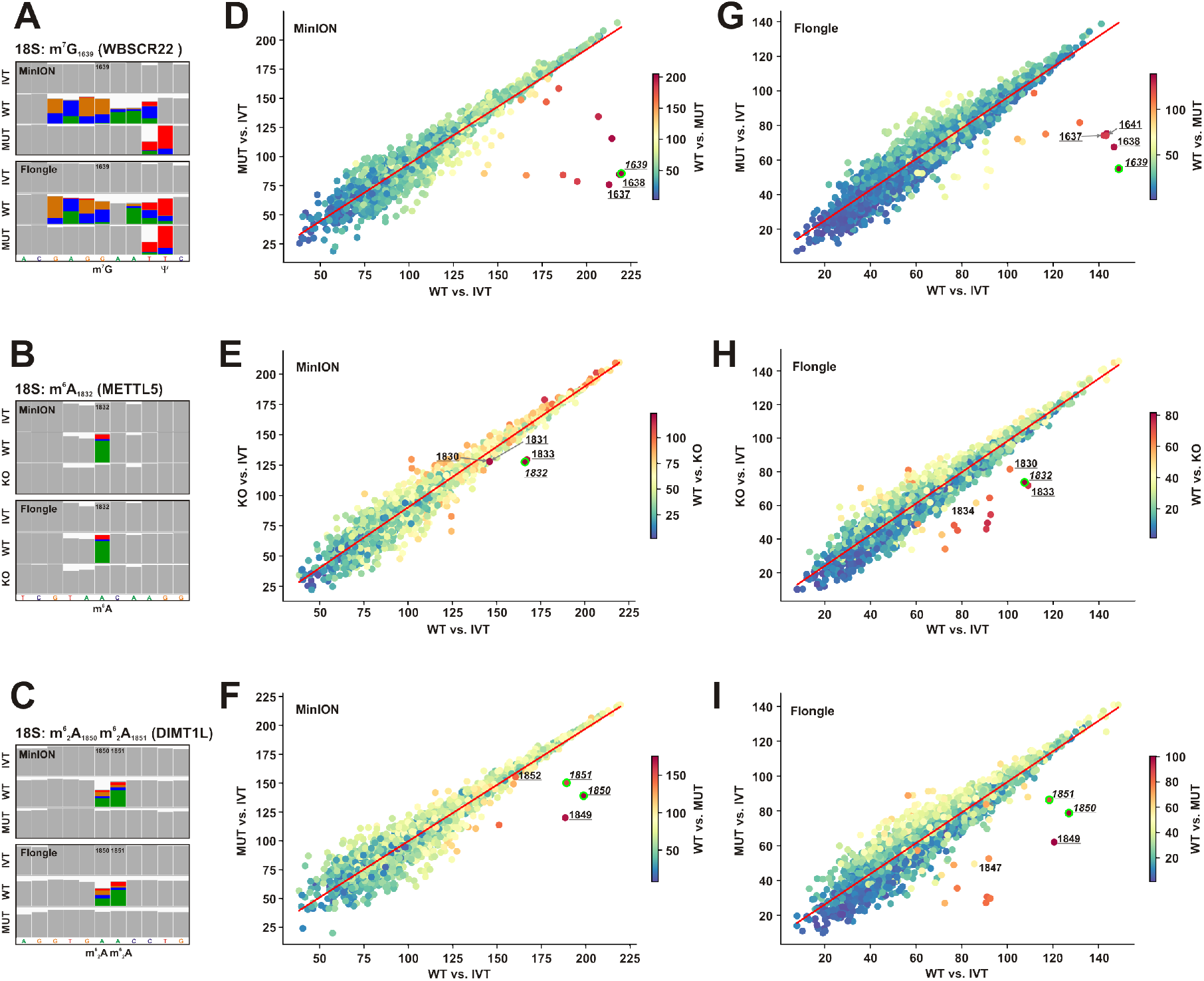
Detection of site-specific modifications in genetically engineered HCT116 cells by Nanopore direct rRNA-seq and JACUSA2 call-2. *In vitro* transcribed 18S rRNA, HCT116 WT and genetically engineered HCT116 cells as listed in Table 1 were subjected to Nanopore direct rRNA-seq, either on a MinION or Flongle flow cell as indicated. A-C) IGV snapshots of the region of interests from MinION or Flongle sequencing as indicated. The target site as well as other described modifications are annotated. Allele frequency threshold = 0.2. D-F) Scatterplot of the three pairwise comparisons of MinION derived data by JACUSA2 call-2 as indicated on the x- and y-axis and in the legend. Labeled are the target site (italics, point indicated by a lime border) as well as significant outliers detected by Local Outlier Factorization (contamination value = 0.002, bold). Outliers in the 5-mer context of the target site are underlined. JACUSA2 features Mismatch Score for 5-mer context, Deletion and Insertion Scores for the site were considered (M_Con_DI). G-I) Scatter plots of the three pairwise comparisons of Flongle derived data as in D-F. A,D,G) Analysis of 18S m^7^G_1639_ in 18S IVT, HCT116 WT and HCT116 WBSCR22^D82K/D82K^. B,E,H) Analysis of 18S m^6^A_1832_ in 18S IVT, HCT116 WT and HCT116 METTL5^-/-^. C,F,I) Analysis of 18S m^6^_2_A_1850_ m^6^_2_A_1851_ in 18S IVT, HCT116 WT and HCT116 DIMT1L^Y131G/Y131G^.

Three different modifications on the 18S rRNA were analyzed in detail: the m^7^G modification at position 1639 catalyzed by WBSCR22 (Fig. 2A,D,G), the m^6^A modification at position 1832 catalyzed by METTL5 (Fig. 2B,E,H) and the double m^6^_2_A modification at positions 1850 and 1851 deposited by DIMT1L (Fig. 2C,F,I). Altered basecalling induced by specific modifications was detected in all analyzed cases as higher Mismatch frequencies in IGV snapshots (Fig. 2A-C, upper panels). For different modifications basecalling was differentially affected. m^7^G_1639_ strongly affects basecalling also of the surrounding residues (−3 to +3), providing a strong signature (Fig. 2A). On the other hand, the m^6^A and m^6^_2_A modifications resulted mainly in Mismatches at the target site (Fig. 2B,C).

We applied JACUSA2 call-2, which compares two different conditions to all possible combinations of the MinION data employing the M_Con_DI Feature set and summarized the results in Scatter plots (Fig. 2D-I). High JACUSA2 scores were obtained for the comparison of WT with either IVT (x-axis) or the mutant cell line (MUT) (colour), but not for the MUT vs. IVT comparison (y-axis) for position 1639 and surrounding residues (Fig. 2D). Statistically significant outliers were calculated with the Local Outlier Factor (LOF) algorithm (contamination value 0.002) (*44*). Strikingly, the target site (lime border, italics) as well as two residues within the 5-mer context were identified as significant outliers (underlined). Similar results were obtained for the two other analyzed modifications (Fig. 2E,F). Importantly, no non-related sites were detected as significant outliers in this 3-way comparison.

Current input requirements may preclude the use of Nanopore direct RNA-seq for this type of samples. Indeed, patient-derived material is usually limited in amounts and analyses have to be carefully planned. To overcome this problem, we aimed to transfer the targeted direct rRNA-seq approach described above to the recently introduced smaller Flongle flow cells (equipped with 126 pores). The overall sequencing results were comparable for MinION and Flongle derived data, as shown exemplarily for HCT116 WT cells (Figure S2A,B). Also the IGV snap-shots for the analyzed modification sites were remarkably comparable between MinION and Flongle data (Fig. 2A-C, lower panels). Importantly, with the 3-way JACUSA2 call-2 analysis we identified all target sites and some neighboring residues, but no non-related sites as significant outliers for all analyzed modifications (Fig. 2G-I).

In summary, all analyzed modifications were detected with JACUSA2 call-2 in the MinION flow cell data as well as in the Flongle flow cell data. In addition, the generated mismatch profile is highly similar between MinION and Flongle sequencing. Overall, the Flongle-based approach enables the analysis of low input samples such as patient derived material making it amenable to clinical biology.

### Nanopore direct rRNA-seq is suitable to estimate modification level

We then determined the required number of reads to detect the investigated modification types. Different read numbers were sampled from the MinION data and subjected to JACUSA2 analysis. As expected, the difference between the JACUSA2 score of target site and the median JACUSA2 score increases, when more reads are considered for analysis, as illustrated for the comparison between WT and KO/MUT (Fig. 3A-D, left panels). A similar result was obtained for the comparison of WT to IVT (Figure S3A-D). The modified sites and / or neighboring residues were consistently detected as significant outliers by 3-way LOF with 300 to 500 (METTL5) reads (Figure S3E-G). To evaluate the robustness of outlier identification, we calculated the normalized distance of the target site LOF scores to the median LOF score (Fig. 3A-D, right panels). Surprisingly, we noticed a small decrease in normalized LOF score distance for WB-SCR22 (Fig. 3A, right panel), which was on the other hand accompanied by a decrease in the standard deviation. Robustness of detection of the METTL5 catalyzed m^6^A modification was increased by higher reads numbers, as indicated by the increased normalized LOF score distance and decreased standard deviation at 5,000 and 10,000 reads, respectively (Fig. 3B, right panel). For the DIMT1L target sites the identification is mostly independent of the number of analyzed reads (Fig. 3C,D).

**Figure 3:**
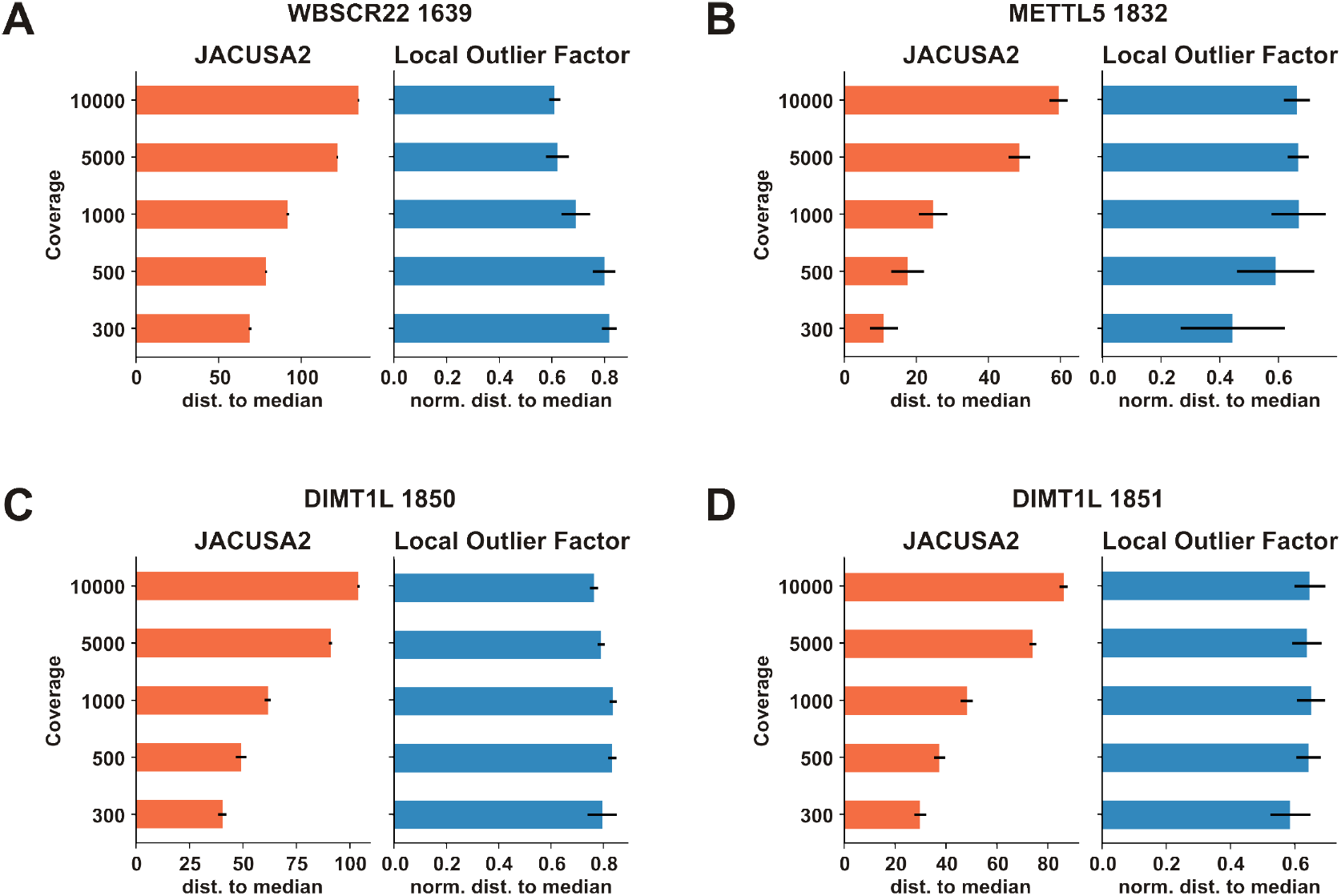
Detection of rRNA modifications with different read numbers. Indicated read numbers were sampled from MinION sequencing data and analyzed by 3-way pairwise comparison as in Figure 2. A) Downsampling analysis of WBSCR22 catalyzed m^7^G_1639_. B) Downsampling analysis of METTL5 catalyzed m^6^G_1832_. C) Downsampling analysis of DIMT1L catalyzed m^6^_2_G_1850_. D) Downsampling analysis of DIMT1L catalyzed m^6^_2_G_1851_. Left panels: Distance of JACUSA2 score for the respective target site to the median JACUSA2 score for the WT versus KO/MUT comparison. Right panels: normalized distance of the 3-way LOF score for the target site to the median LOF score. Shown are the mean and the standard deviation from down sampling employing different seeds (n = 15).

The down sampling analysis indicated that 1,000 reads are sufficient to detect the analyzed modifications when comparing completely modified reads (WT) with non-modified reads (IVT, KO/MUT). However, in physiological or pathological settings, more subtle changes in modification level are expected. We therefore analyzed to what extent rRNA modifications can be analyzed at substoichiometric levels as well. To do so, we sampled a total of 1,000 reads from all samples. For the reference “WT” sample, different ratios of WT and KO/MUT reads were bioinformatically mixed as indicated (Fig. 4). Samples from all mixing ratios derived from 5 seeds were analyzed by the 3-way JACUSA2 comparison as outlined above. Interestingly, m^7^G_1639_ and m^6^_2_A were consistently identified with only 5% modified reads (Figure S4E,G,H) based on the target site and/ or neighboring residues. m^6^A had a detection threshold around 25% (Figure S4F). Importantly, an increase in the JACUSA2 score with increasing modification frequency was detected for all analyzed modification sites (Fig. 4), indicating that Nanopore direct rRNA-seq can also be used for estimation of modification levels. For m^6^A_1832_, a linear relation of JACUSA2 score and modification level was observed for the comparison to both, KO and IVT (Fig. 4B). For the other modification types, the JACUSA2 score seems to approach saturation (Fig. 4A,C,D). To exclude an effect of the seed used for downsampling, the analysis was repeated with a different seed, which yielded comparable results (Figure S4A-D). In conclusion, our *in silico* mixing approach shows that rRNA modification levels can be estimated from Nanopore direct RNA-seq data based on appropriate calibration curves.

**Figure 4:**
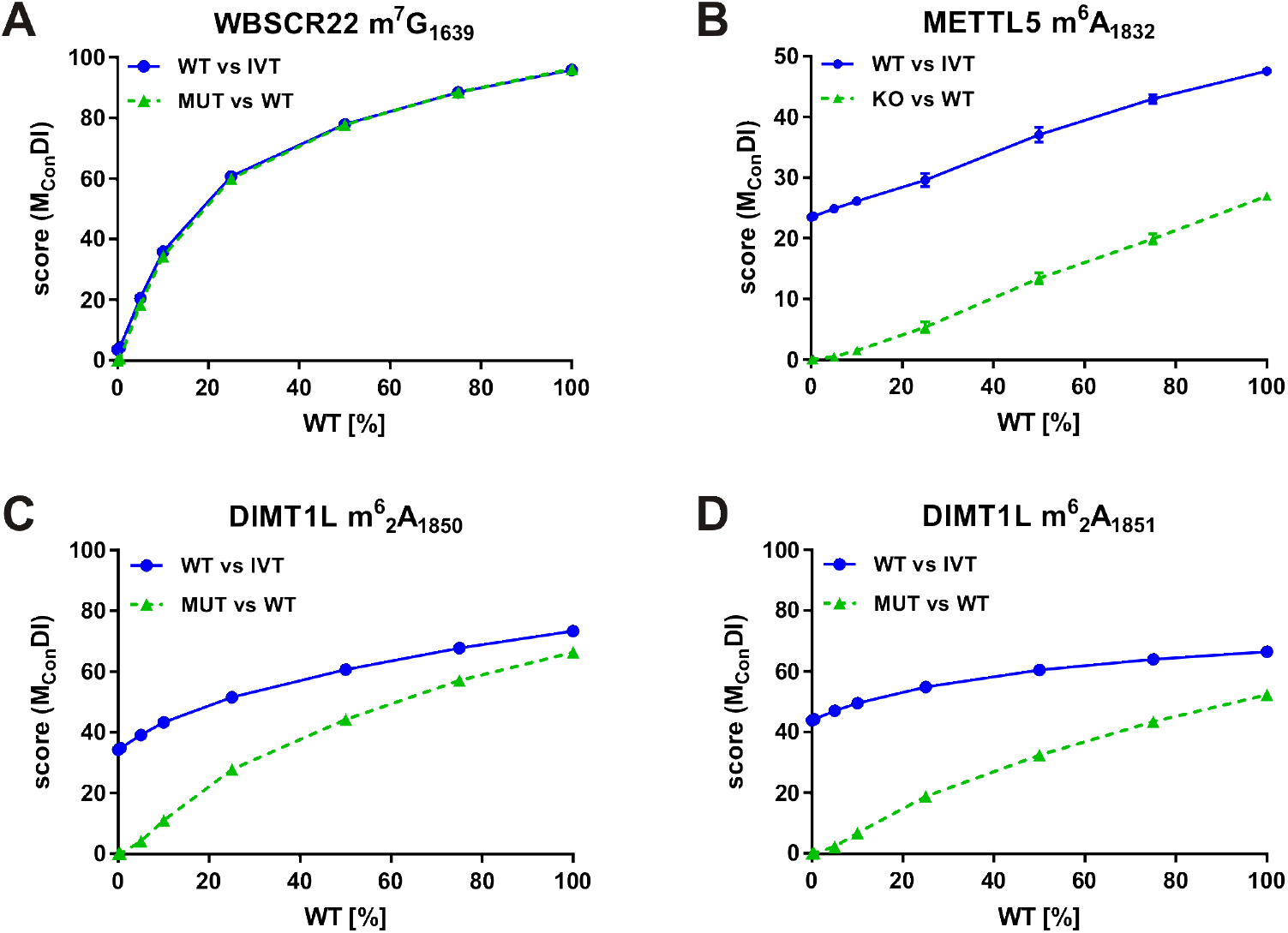
Analysis of the influence of modification level on the JACUS2A score. 1,000 reads were downsampled with seed “0” from the MinION sequencing data shown in Figure 2. The “WT” sample was composed of modified (WT) and unmodified (KO/MUT) reads as indicated that were derived from the downsampled data with 5 different seeds. JACUSA2 call-2 analysis considering the M_Con_DI Feature set. The modification rate vs. the JACUSA2 score of the respective target site is shown (n =5). A) Analysis of m^7^G_1639_. B) Analysis of m^6^A_1832_. C) Analysis of m^6^_2_A_1850_. D) Analysis of m^6^_2_A_1851_.

### rRNA modifications are dynamic during cardiac differentiation of human induced pluripotent stem cells

The comparative analysis of Nanopore direct rRNA-seq data allows the detection of changes in specific modifications such as base methylations (Fig. 2). Furthermore, we have shown that JACUSA2 can be applied to classify non-modified and modified uridine residues in Illumina and Nanopore direct RNA data (*34*). We reasoned that JACUSA2 combined with Nanopore direct rRNA-seq is able to identify any changes in rRNA modifications that are detectable by their basecalling signature (Mismatch, Deletion and Insertion score). The comparative analysis of different cellular states, such as during cell differentiation, facilitates the handling of unbalanced classes of unmodified and modified sites, especially for infrequent base modifications.

To test this hypothesis, we analyzed rRNA from human induced pluripotent stem cells (hiPSC) during their differentiation to cardiomyocytes. With the applied protocol, cells usually start beating at day 9, indicating their successful differentiation to cardiomyocytes. Therefore, we compared rRNA modifications at day 0, day 5 and day 9 of differentiation (Fig. 5). Samples from two independent differentiation experiments were sequenced to a depth of about 50,000 raw reads on MinION flow cells and treated as replicates in JACUSA2. As rRNA modifications are clustered in functional relevant regions and we did not had *a priori* knowledge on expected changes, we chose the MDI Feature set here in order to prevent confounding of the analysis by neighboring sites. Furthermore, we applied very stringent conditions for outlier detection with a LOF contamination value of 0.001 to focus only on highly confident sites. With these parameters, we identified 18S Um_354_ and three modified sites on the 28S rRNA (psU_4493_, Gm_4494_, psU_4673_; 28S numbering refers to (*6,45*)) to be significantly changed during cardiomyocyte differentiation (Fig. 5A,C). All changes were also significant in the comparison of day 0 versus day 9 (Fig. 5B,D), whereas from the other pairwise comparisons alone only 28S psU_4673_ (Figure S5A-D) was identified. The representative IGV snapshots indicate an increase in Um_354_ modification in the course of the differentiation (Fig. 5E, upper panel), which is further supported by the gradual increase of U-to-C transitions and deletions during differentiation (Fig.5E, lower panel). This 2’-O-methylation site was previously identified (*8, 46*) and reported to be 20% methylated (*8*) in the human lymphoblast cell line TK6. Interestingly, Um_354_ was described to be upregulated during development of zebrafish (*47*) and mouse brain (*48*). Similar to 18S Um_354_, also 28S U_4673_ was found to be hypomodified (*8*) (39%) and the pseudouridylation increases during cardiomyocyte differentiation as well (Fig. 5F). The psU and Gm modifications at 28S 4493/4494 were reported to be 17% and 91% present in TK6 cells, respectively (*8*) and are enhanced during cardiomyocyte differentiation (Fig. 5G). For all differentially modified sites, no specific function in ribosome biogenesis or mRNA translation has been assigned to this day.

**Figure 5:**
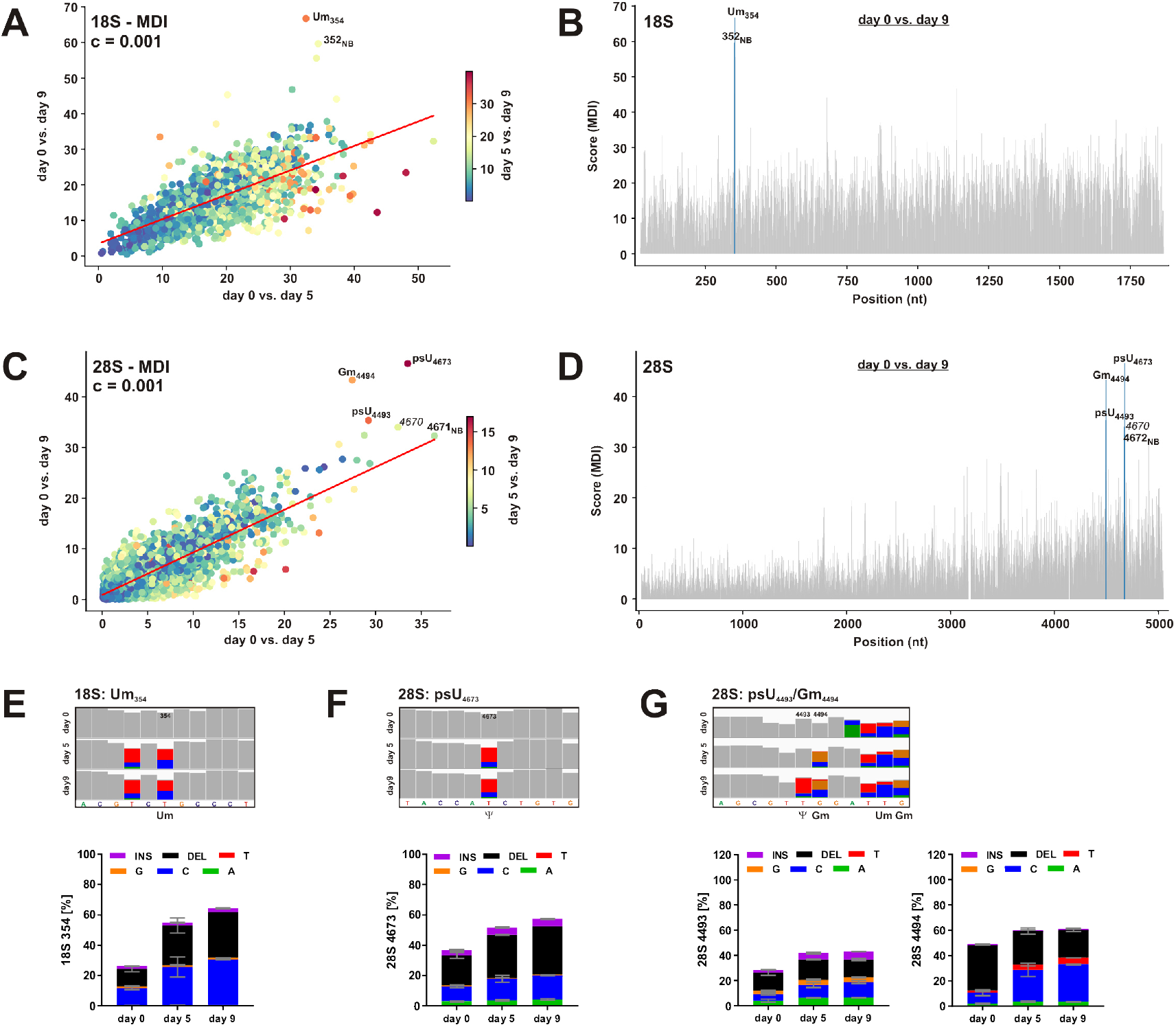
Analysis of rRNA modifications changes in cardiomyocyte differentiation. Human induced pluripotent stem cells were differentiated to cardiomyocytes for up to 9 days in two independent replicates. RNA was isolated on days 0, 5 and 9 and subjected to Nanopore direct rRNA-seq on MinION R9.4.1 flow cells. Runs were stopped after approximately 50 K reads were obtained. A) Scatter plot of JACUSA2 call-2 analysis of all possible pairwise comparisons for 18S rRNA. The MDI Feature set was applied. Significant outliers detected by 3-way LOF (contamination value 0.001) are labeled. B) Bar plot of the pairwise comparison day 0 vs. day 9 for 18S rRNA. Significant outliers detected by LOF (contamination value 0.001) are labeled. C) Scatter plot of JACUSA2 call-2 analysis of all possible pairwise comparisons for 28S rRNA. The MDI Feature set was applied. Significant outliers detected by 3-way LOF (contamination value 0.001) are labeled. D) Bar plot of the pairwise comparison day 0 vs. day 9 for 28S rRNA. Significant outliers detected by LOF (contamination value 0.001) are labeled. E) Upper panel: IGV snapshot of 18S Um_354_ and surrounding residues. Allele frequency threshold = 0.2. Lower panel: Basecalling errors at position 18S U_354_ during cardiomyocyte differentiation. F) Upper panel: IGV snapshot of 28S psU_4673_ and surrounding residues. Allele frequency threshold = 0.2. Lower panel: Basecalling errors at position 28SU_4673_ during cardiomyocyte differentiation. G) Upper panel: IGV snapshot of 28S psU_4493_, Gm_4494_ and surrounding residues. Allele frequency threshold = 0.2. Lower panel: Basecalling errors at positions 28S U_4493_ (left) and G_4494_ (right) during cardiomyocyte differentiation.

The analysis of rRNA modifications in hiPSC-derived cardiomyocyte differentiation shows that Nanopore direct rRNA-seq is suitable to detect changes in RNA modification pattern with limited amount of sample and can detect a wide range of rRNA modifications: 2’-O-methylation of ribose, isomerization of uridine to pseudouridine, as well as other base modifications (Fig. 2).

### Identification of rRNA modification changes in cardiac biopsies

The analysis of hiPSC-derived cardiomyocyte differentiation revealed that Nanopore direct rRNA-seq in combination with JACUSA2 detects dynamic changes in rRNA modifications. Furthermore, the in-depth analysis of the HCT116 genetic model system and respective IVTs has shown that the output of Flongle sequencing is sufficient to identify changes in rRNA modifications. Importantly, this makes direct rRNA-seq accessible to the analysis of patient-derived samples, enabling the analysis of so far underexplored epitranscriptomics in human diseases.

To directly illustrate this, we analyzed biopsy samples from patients suffering from dilated cardiomyopathy (DCM) or hypertrophic cardiomyopathy (HCM) and compared their modification pattern to samples from heart-transplanted control patients (HTX). Two independent samples from each condition were analyzed by JACUSA2 as replicates to identify consistent changes for each disease. To prevent disturbance of the analysis by neighboring sites, we applied the MDI Feature set without 5-mer context and a LOF contamination value of 0.001, as previously used for the analysis of cardiomyocyte differentiation. Interestingly, we identified ac^4^C_1842_ as significantly different site on the 18S rRNA for both, the DCM and HCM samples compared to the control samples (Fig. 6A,C). Furthermore, modification of psU_4299_ on the 28S rRNA wass significantly different in DCM but not HCM patients compared to the control samples (Fig. 6B,D). The acetylation of C_1842_ seems to be increased in patient samples compared to the HTX control (Fig. 6E,F). This acetylation is catalyzed by NAT10 (*49,50*), which is essential for pre-18S rRNA processing. Compared to the neighboring residues, psU_4299_ shows low coverage in all analyzed samples, however an especially increased deletion rate in DCM indicates a higher modification level under disease conditions. Interestingly, differences in psU_4299_ were not found in the HCM samples (Fig. 6G,H).

**Figure 6:**
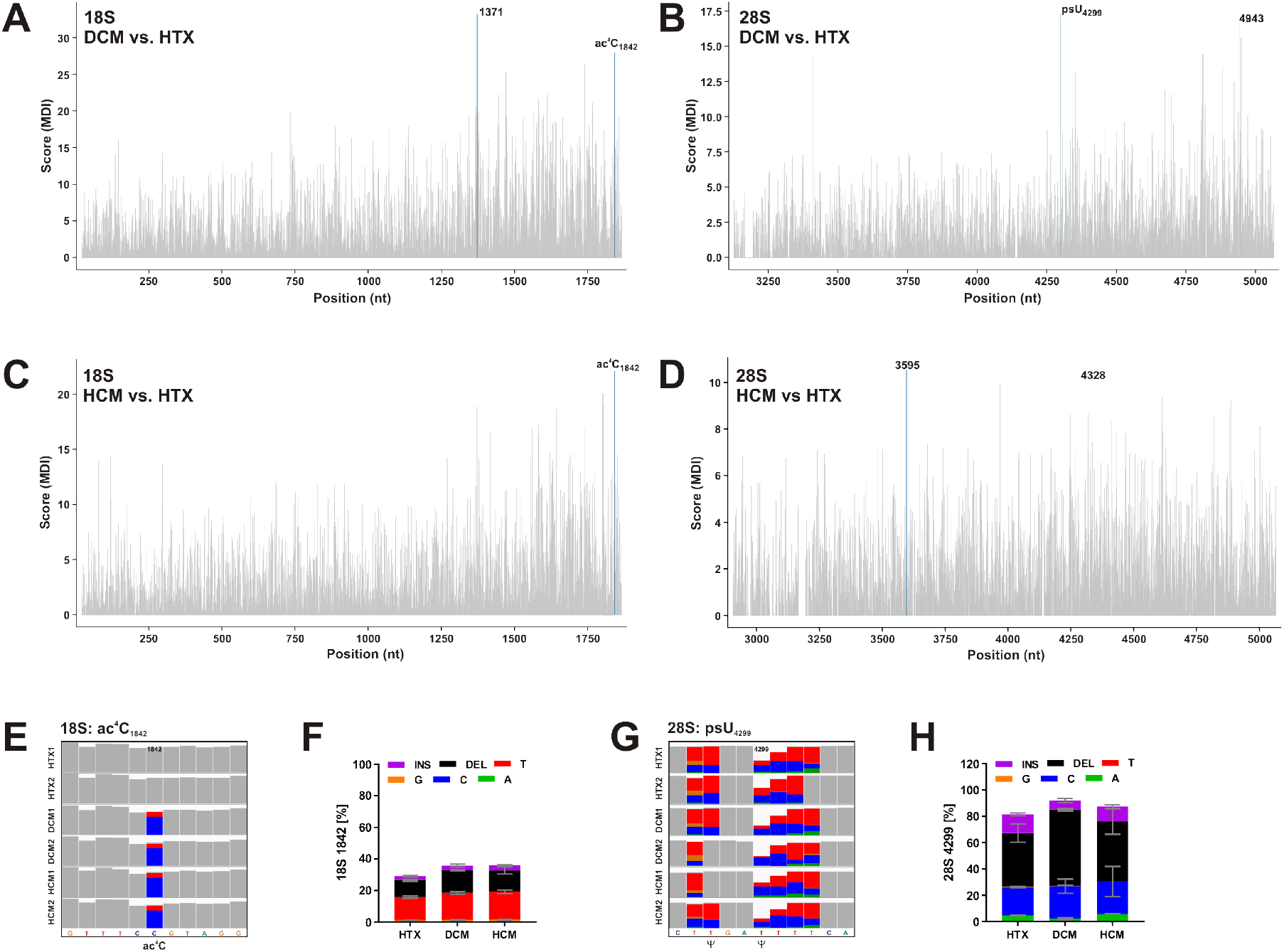
Detection of changes in rRNA modifications by Nanopore direct rRNA-seq in samples from patients with cardiomyopathy. Nanopore direct rRNA-seq was applied to total RNA isolated from biopsies of two heart transplanted patients (HTX, ctrl.), two DCM and two HCM patients, respectively. The two samples from each condition were analyzed as replicate samples with JACUSA2 call-2. A) Barplot of JACUSA2 scores for 18S rRNA for the DCM vs. HTX comparison considering the MDI Feature set. Significant outliers detected by LOF (contamination value = 0.001) are labeled and highlighted in blue. B) Analysis of 28S rRNA as in A. C) Barplot of JACUSA2 scores for 18S rRNA for the HCM vs. HTX comparison considering the MDI Feature set. Significant outliers detected by LOF (contamination value = 0.001) are labeled and highlighted in blue. D) Analysis of 28S rRNA as in C. E) IGV snapshot of 18S position C_1842_ and surrounding residues. Allele frequency threshold = 0.2. F) Basecalling errors at position 18S C_1842_. G) IGV snapshot of 28S position U_4299_ and surrounding residues. Allele frequency threshold = 0.2. H) Basecalling errors at position 28S C_4299_.

Additionally, we identified a few significant outliers that, to our knowledge, are not related to any known modification site (18S U1371, 28S U3595, G4328 and A4943). These positions may be caused by a higher noise in this analysis due to lower quality of the analyzed RNA or, possibly, may reflect differential expression of rRNA variants.

In this study, we establish the Nanopore-based targeted direct RNA-sequencing of human ribosomal RNAs and analyze widespread rRNA modifications employing JACUSA2. We show that direct rRNA-seq on the Nanopore can be scaled down to the Flongle device, enabling the analysis of patient derived material. The analysis of patient derived heart biopsies revealed possible disease associated changes in modification of 18S ac^4^C_1842_ and 28S psU_4299_.

## Discussion

Modifications of human rRNAs are well studied, but until recently considered as relatively static, as the complete ribosome. Emerging evidence however supports the hypothesis of ribosomal heterogeneity (*51*) that may contribute to the etiology and progression of several diseases, including cancer and cardiovascular diseases. Ribosomal heterogeneity can arise by different means: use of alternative ribosomal protein paralogs, expression of different rRNA variants or differential modifications of rRNAs and ribosomal proteins. Furthermore, mutations in ribosomal components or ribosome assembly factors may lead to a disease class summarized as “ribosomopathies” (*15*). Ribosomopathies are mostly developmental diseases manifesting a large spectrum of tissue-specific defects. One classical example is Diamond-Blackfan anemia, which mainly affects maturation of red blood cells (*16*). While mutations in ribosomal proteins are studied relatively well, only few studies analyzed differences in variants or differential modifications of rRNAs independent of the modification type. The analysis of rRNA mutations has been complicated by the complex genomic organization of the rDNA loci that are located in tandem repeats on several chromosomes (*26*). Long read sequencing techniques should fill this gap of the human genome project soon (*52*).

The sequence specific and quantitative analysis of various RNA modification types with moderate amounts of input material is just currently evolving based on the direct RNA-sequencing method. We and others have shown recently that Nanopore direct RNA-seq in combination with suitable computational analysis frameworks can be used to detect different RNA modifications (*31, 33-36, 38*). However, current analyses focused mostly on specific types of modifications (m^6^A, pseudouridine) or did not address dynamic changes in human samples with clinical relevance.

Here we established the targeted direct RNA-seq of human ribosomal RNA with custom adapters for 18S and 28S rRNA (direct rRNA-seq). We observed a lower efficiency of 28S rRNA sequencing throughout our analyses with about tenfold less and also shorter reads from 28S compared to 18S rRNA (Fig. 1E). This is most likely not the consequence of a lower efficiency of the 28S adapter, but caused by the high GC content in parts of the 28S rRNA (more than 80%) and the long homopolymeric stretches, which consistently were previously reported to disturb Nanopore sequencing (*42, 43*).

We have used direct rRNA-seq for a comprehensive analysis of base modifications on human rRNA based on genetic model systems and IVTs, and show that the recently introduced JACUSA2 framework (*34*) detects all analyzed 18S rRNA modifications in data from the standard MinION flow cells, namely m^7^G_1639_, m^6^A_1832_, and the double di-methylation m^6^_2_A_1850_ m^6^_2_A_1851_. The different base modifications cause distinct basecalling errors that are reflected in the Mismatch, Deletion and Insertion scores calculated by JACUSA2 (Fig. 2). We show that different Feature sets may be used to detect differences in rRNA modification (Figure S1). Whereas the consideration of the 5-mer context causes high signal-to-noise ratio in relatively low modified regions, the context may confound analysis in a highly modified context. In such circumstances, a dedicated Feature set focused on the analyzed site may be more appropriate. We found the Local Outlier Factorization (LOF) (*44*) for detection of significant different sites superior compared to a residual based method (not shown). This may imply that the modification signal yields a high level of skewness within local regions. Sensitivity and specificity of this method may be adjusted by the contamination value as well as the number of considered neighbors. The outlierness can be measured based on the density of regions of twenty points and, consequently, the modified positions are constantly the top ranked outliers.

Importantly, we have reported that the detection of rRNA modifications can be scaled down to the smaller Flongle flow cells (Fig. 2), enabling the analysis of patient derived material available only in limited amounts (Fig. 6). This is of special interest, as barcoding of direct RNA-seq libraries is currently not officially supported by ONT. Sequencing on the Flongle flow cells yields data with comparable quality, and detection of modifications is only dependent on the number of obtained reads that is determined by the quality of the Flongle flow cells. By downsampling analyses from the MinION data, we show that 300 to 500 reads are sufficient to identify the analyzed modification sites (Fig. 3). Interestingly, although the JACUSA2 scores increase with read number, the identification of the sites as outliers by LOF is highly robust in most cases (Fig. 3). Only for the METTL5 catalyzed m^6^A modification, higher reads number were found to be beneficial.

Importantly, employing *in silico* mixing of modified and non-modified reads, we show that the modification level of selected base modifications can be estimated from the JACUSA2 scores based on a calibration curve (Fig. 4). JACUSA2 was previously described to accurately identify modified uridine residues (pseudouridine and 2’-O-methylated uridine) in direct rRNA-seq data (*34*). Although we did not cover the complete repertoire of rRNA modifications at the time, our data indicate that applying JACUSA2 - direct rRNA-seq is able to capture all RNA modifications that are detectable by basecalling errors (Mismatch, Insertion, Deletion).

Our main goal was to establish the analysis of dynamic changes in rRNA modifications on low input samples. It was reported previously in mouse and zebrafish that rRNA modifications change during development and differentiation (*47, 48*). We therefore analyzed Nanopore direct rRNA-seq data from two replicates of hiPSC-cardiomyocyte differentiation for changes in rRNA modifications (Fig. 5). Indeed, we identified high confident sites on the 18S and 28S rRNA as differentially modified, namely an increase in modification of 18S Um_354_ and several 28S sites (psU_4493_, Gm_4494_, psU_4673_) (Fig. 5) was detected. Interestingly, an increase of Um_354_ has been reported previously during development of zebrafish and mouse brain as well (*47,48*). This analysis shows that dynamic changes in rRNA modifications can be studied with Nanopore direct RNA-seq. Importantly, identification of these changes is not dependent on IVT and genetic model systems, which may not be available in most cases.

Finally, we analyzed a set of human heart biopsies for differences in rRNA modifications. We made use of the JACUSA2 option to handle replicates and treated the two analyzed independent patient samples for each condition as such. In this analysis we detected an increase in ac^4^C_1842_ in DCM and HCM patients compared to control patients (HTX) (Fig. 6). Furthermore, differential modification of psU_4299_ was observed for DCM, but not HCM patients. These data show that Nanopore direct rRNA-seq - JACUSA2 enables the *de novo* identification of disease associated changes in rRNA modifications.

In this work, we apply JACUSA2 call-2 in pairwise comparisons to identify differences in rRNA modification. A number of other computational frameworks have been published to analyze rRNA modifications in Nanopore direct RNA-seq data including EpiNano (*37*), xPore (*36*) and ELIGOS (*33*), which are either based on the detection of basecalling errors as JACUSA2 or derive modification pattern from changes in the raw data. The major challenge in the analysis of RNA modifications compared to DNA modifications is the diversity of modification types. In contrast to most other frameworks, JACUSA2 supports the handling of replicate samples as well as pairwise comparison (*34*), enabling either the comparison to a non-modified reference sequence or the identification of differences between biological samples. Furthermore, we found JACUSA2 to be superior in terms of analysis time.

We established the targeted Nanopore direct RNA-seq of human rRNA and detection of modifications applying JACUSA2. The down scaling to Flongle flow cells enabled the study of patient-derived samples as human heart biopsies and revealed several candidate site for differential modification during cardiac differentiation and pathogenesis. These analyses show that differential rRNA modification should be considered in the analysis of disease pathogenesis and development. Further work will be required to understand the biological consequences of these altered modification patterns.

## Methods

### Generation of HCT116 mutant cell lines

All human cell lines were generated in p53-positive diploid HCT116 cells (ATCC, #CCL-247) by genome editing. The recipient cell line was diagnosed by ATCC by short tandem repeat (STR) analysis, prior to use. The HCT116 METTL5^-/-^ cell lines has been described previously (*39*). Here, the exon encoding the catalytic domain of the protein was precisely excised from the genome on both alleles by CRISPR-Cas9 genome editing.

The HCT116 DIMT1L^Y131G/Y131G^ and WBSCR22^D82A/D82A^ will be described in detail elsewhere. Briefly, the selected point mutation was introduced by CRISPR-Cas9 genome editing on both alleles, as follows: an *in vitro* reconstituted Cas9 RNP complex (final concentration 4 μM) consisting of a specific crRNA guide (see Table S1), a universal tracRNA (IDT, #1072532), and the Streptococcus pyogenes Cas9 (IDT, #1081058), was electroporated in cells freshly resuspended in nucleofector solution V (Lonza, VCA-1003) together with a single-strand donor DNA (ssDNA, final concentration 4 μM) carrying the mutation. Cells were electroporated with enhancer (IDT, #1075915, final concentration 4 μM) in a nucleofector device (Lonza, Nucleofector 2; program D-032). Cells were incubated for 24 h to allow them to recover and then detached and cloned by serial dilution. Individual clones were selected, diagnosed by differential restriction digest of a PCR-amplified product of the modified area (gain of a *Bst*NI restriction site for the DIMT1L^Y131G/Y131G^ clones, or loss of a *Eco*RV restriction site for the WBSCR22^D82A/D82A^ clones), by DNA sequencing of the modified area, and by loss of 18S rRNA modification by primer extension as described (*40, 53*).

### Cell culture

HCT116 cells were cultured in McCoy’s 5A Medium (Lonza, BE12-168F) supplemented with 10% fetal bovine serum (Sigma, F7524) and 100 U/ml penicillin and 100 μg/ml streptomycin (Lonza, DE17-602E) in a New Brunswick Galaxy 170R incubator at 37°C and under 5% CO_2_.

WT1.14 cells were a kind gift from S. Doroudgar and differentiated based on described protocols (*54,55*). Briefly, differentiation was started with 80-100% confluent hiPSC cultures by addition of Cardio Differentiation medium (RPMI1640 with HEPES and GlutaMax, 0.5 mg/ml human recombinant albumin, 0.2 mg/ml L-ascorbic acid 2-phosphate) supplemented with 4 μM CHIR99021. After 24 h, medium was exchanged to Cardio Differentiation medium. On day 2, cells were cultured in Cardio Differentiation Medium supplemented with 5 μM IWP2 for two days and afterwards transferred to Cardio Differentiation Medium for another two days. On day 7, medium was changed to Cardio Culture medium (RPMI1640 with HEPES and GlutaMax, supplemented with B27 supplement with Insulin). RNA was isolated on day 0, 5 and 9 of cardiac differentiation.

### Patient samples

Heart biopsies from patients diagnosed with dilated cardiomyopathy (DCM) or hypertrophic cardiomyopathy (HCM) were obtained from the Heidelberg Cardio Biobank (HCB) approved by the ethics committee (Application No. S-390/2011). Samples from heart transplanted (HTX) patients obtained from the HCB were used as control. The study was conducted according to the principles outlined in the Declaration of Helsinki. All participants have given written informed consent to allow for molecular analysis. Only samples from male patients with an age of 50 to 65 years were included. Subject to availability either one or two biopsies were used for isolation of total RNA as described below after homogenization.

### Isolation of total RNA

Total RNA from HCT116 cells was extracted in TriReagent solution (Thermo Fisher) according to the manufacturers instructions.

Total RNA from WT1.14 cells was isolated using Trizol. Cells were washed with ice-cold PBS, scraped in 750 μl Trizol and incubated 5 min at room temperature before addition of 200 μl chloroform. Samples were centrifuged (20 min, 10,000 g, room temperature) and the aqueous phase re-extracted with one volume chloroform: isoamylalkohol (24:1) (5 min, 10,000 g, room temperature). The RNA in the aqueous phase was precipitated with one volume isopropanol (30 min, 20,800 g, 4°C), washed twice with 1ml 80% ethanol in DEPC-H_2_O and dissolved in 25 μl DEPC-H_2_O (10 min, 55°C, shaking).

### Isolation of genomic DNA

Genomic DNA was isolated from 5 Mio HeLa cells using the NucleoSpin tissue kit (Macherey-Nagel) according to the manufacturers protocol.

### Generation of templates for *in vitro* transcription

The complete 18S rRNA sequence and the 3’ fragment of 28S rRNA (F3, nts 3619 to 5070) were amplified from genomic DNA by touchdown PCR with Q5 DNA polymerase (New England Biolabs) using a forward primer that introduces the T7 promoter sequence for *in vitro* transcription (IVT). The following protocol was used for touchdown PCR: 30 sec initial denaturation at 98°C, 20 cycles of touchdown (10 sec, 98°C; 20 sec, 72°C to 62°C (ΔT_m_ −0.5 °C); 5 min, 72°C), followed by 15 cycles standard PCR at 62°C annealing temperature and final elongation (5 min, 72°C). Primers sequences were: T7-18S fw: TAATACGACTCACTATAGtacctggttgatcctgccag, 18S rv: taatgatccttccgcaggttc, T7-28S-F3 fw: TAATACGACTCACTATAGggaccaggggaatccgactg, 28S rv: gacaaacccttgtgtcgaggg.

### *In vitro* transcription

IVTs were generated using T7 Megascript kit (Thermo Fisher Scientific) according to the manufacturers protocol. RNA integrity was analyzed on a 1% agarose gel. IVT products were purified using RNA Clean and Concentrator kit (Zymo Research).

### Polyadenylation of 18S IVT

1 μg 18S IVT was polyadenylated with a E-PAP based Poly(A) Tailing Kit (Thermo Fisher Scientific) according to the manufacturers instructions and purified using RNA Clean and Concentrator kit (Zymo Research).

### Generation of ONT direct RNA seq libraries for sequencing on FLO-MIN106D (R9.4.1) flow cells

Direct RNA-seq libraries were generated using the SQK-RNA002 kit (Oxford Nanopore Technologies) following the sequence-specific protocol. Universal oligo A and sequence-specific oligo B were annealed at a concentration of 1.4 μM each in 10 mM Tris, pH 7.5, 50 mM NaCl (2 min, 95°C; 0.1°C/sec to 22°C). Briefly, 500 ng total RNA in a volume of 9 μl was ligated to 1 μl custom adapter mix (18S:28S ratio 1:2) using 1.5 μl T4 DNA ligase (New England Biolabs) in NEB next Quick ligation buffer (3 μl, New England Biolabs) in the presence of 1 μl RNA CS (Oxford Nanopore Technologies) for 10 min at room temperature. Reverse transcription to stabilize the RNA strand was performed using Superscript IV reverse transcriptase (Thermo Fisher Scientific for 50 min at 50°C, followed by enzyme inactivation (10 min, 70°C). Reactions were cleaned up using Agencourt RNAClean XP beads (Beckman Coulter). The RMX RNA adapter was ligated as described above, followed by Agencourt RNAClean XP purification and elution in 21 μl elution buffer. The concentration of the library was determined using Qubit DNA HS assay (Thermo Fisher Scientific). The sequences of the adapters were: oligo A: /5PHOS/GGCTTCTTCTTGCTCTTAGGTAGTAGGTTC, 18S v1 oligo B: GAGGCGAGCGGTCAATTTTCCTAAGAGCAAGAAGAAGCCtaatgatcct, 18S v2 oligo B: GAGGCGAGCGGTCAATTTTCCTAAGAGCAAGAAGAAGCCtaatgatccttccgcaggtt, 28S oligo B: GAGGCGAGCGGTCAATTTTCCTAAGAGCAAGAAGAAGCCgacaaaccct. Libraries were sequenced on a GridION X5 device equipped with MinION R9.4.1 flow cells for 48 h in high-accuracy mode.

Libraries from cardiomyocyte differentiation samples were prepared with 200 ng according to the above protocol. Sequencing runs were stopped after approximately 50 K reads were obtained.

### Generation of ONT direct RNA seq libraries for sequencing on Flongle flow cells

Libraries for sequencing on Flongle flow cells were prepared as above with some modifications. Libraries were prepared with 200 ng total RNA as input. Ligation steps were extended from to 15 min, incubation during beads purification to 10 min. After the first cleanup, RNA was eluted in 10 μl H_2_O and the following steps carried out in a smaller volume: 10 μl RNA were ligated to 2 μl RMX with 1 μl T4 DNA Ligase in a total volume of 20 μl and purified with an equal volume of Agencourt RNAClean XP beads. Libraries were eluted in 9 μl ELB. Flongle flow cells were loaded by a community protocol to allow loading similar to the FLO-MIN106D flow cells (https://community.nanoporetech.com/posts/a-very-gentle-relatively). Flongle flow cells were primed with 117 μl FLB + 3 μl FLT. 8 μl library was diluted with 7 μl H_2_O and loaded with 15 μl RRB. Flongle libraries were sequenced on a GridION X5 device equipped with Flongle adapters and Flongle flow cells for 24 h in high-accuracy mode.

### Preprocessing of direct RNA-seq data

Reads were mapped using minimap2 version 2.17. BAM files were filtered to exclude secondary and poor alignments. Plus, the MD tag was added to allow variant calling using JACUSA2 software without the need for the reference transcriptome, then, the resulting BAM files were indexed.

### Detection of modifications with JACUSA2

JACUSA2 software was used to generate differential features summarizing essentially the Mismatch, Deletion, and Insertion scores for variant discrimination, by comparing different conditions using the call-2 option and integrating information from replicate experiments where applicable. The Feature estimation was followed by modification detection using different Feature sets. Only residues with a coverage of more than 30 reads in all conditions and replicates were considered in the analysis. Local Outlier Factor (LOF) (*44*) method was used to detect the modifications. LOF, in principle, predicts outliers in an unsupervised manner by measuring the density deviation of each point with respect to its neighbors. Hence, it assigns for each point a score of being outlying, then, the ensemble of the most outlying points (the highest scores) is identified. The proportion of outliers to be captured for the analysis cases was set to 0.1-0.2% (contamination value 0.001-0.002) and neighborhood size was 20. Plots were built using matplotlib.

### Downsampling analysis

To evaluate the effect of read coverage on the analysis, BAM files were downsampled to different amounts of reads (0.3k, 0.5k, 1k, 5k,10k). To ensure the randomness of read selection, various seed values were considered. The generated down-samplings were subjected to the 3-way JACUSA2 call-2 analysis. To compare results across the different levels of read coverage, we calculated the distance between modification sites and the median in terms of two basic scores: the combined feature M_Con_DI from the estimated JACUSA2 call-2 scores and the score assigned to each site by LOF method. To avoid bias caused by the different scales of LOF scores across analyses, the normalized distance was considered, so that the difference between the score of the modification site and the median is divided by the maximum LOF value.

### *In silico* mixing analysis

To evaluate the ability to detect rRNA modifications with low stoichiometry, *In silico* samples with different average of modification rates (0%, 0.5%, 5%, 10%, 25%, 50%, 75%, 100%) were designed by combining WT and MUT samples of 1k reads derived from two different seed values (“0” and “42”). Then, differential analysis of the generated mixtures and the MUT/ IVT samples was performed using JACUSA2 call-2. The Feature set M_Con_DI from the estimated JACUSA2 scores was compared across the different mixture ratios.

Preprocessing, down-sampling, and MUT/WT mixing were performed using Samtools version 1.9. A Snakemake pipeline of the analysis workflow was developed and is available on Zenodo (DOI: 10.5281/zenodo.5654450).

## Acknowledgments

This work was supported by the Klaus Tschira Foundation. Research in the lab of DLJL is supported by the Belgian Fonds de la Recherche Scientifique (F.R.S./FNRS), the Université Libre de Bruxelles (ULB), the European Joint Programme on Rare Diseases (‘RiboEurope’ and ‘DBAcure’), the Région Wallonne (SPW EER) (‘RIBOcancer’), and the Internationale Brachet Stiftung.

## Author contributions

CD supervised the project. CD, DLJL and ISNdV designed the study. ISNdV performed all experiments except otherwise stated. CZ and DLJL generated HCT116 cell lines and provided RNA. MB and JE performed hiPSC-CM differentiation experiments. JE and SS supported Nanopore sequencing. SS and BM identified and providied heart biopsy samples. AL, ISNdV and CD analyzed data and contributed figures. ISNdV wrote the manuscript. All authors critically revised the manuscript.

## Supplementary materials

Supplementary Information

Figs. S1 to S5

## Supplementary Information

**Supplementary Table 1:**
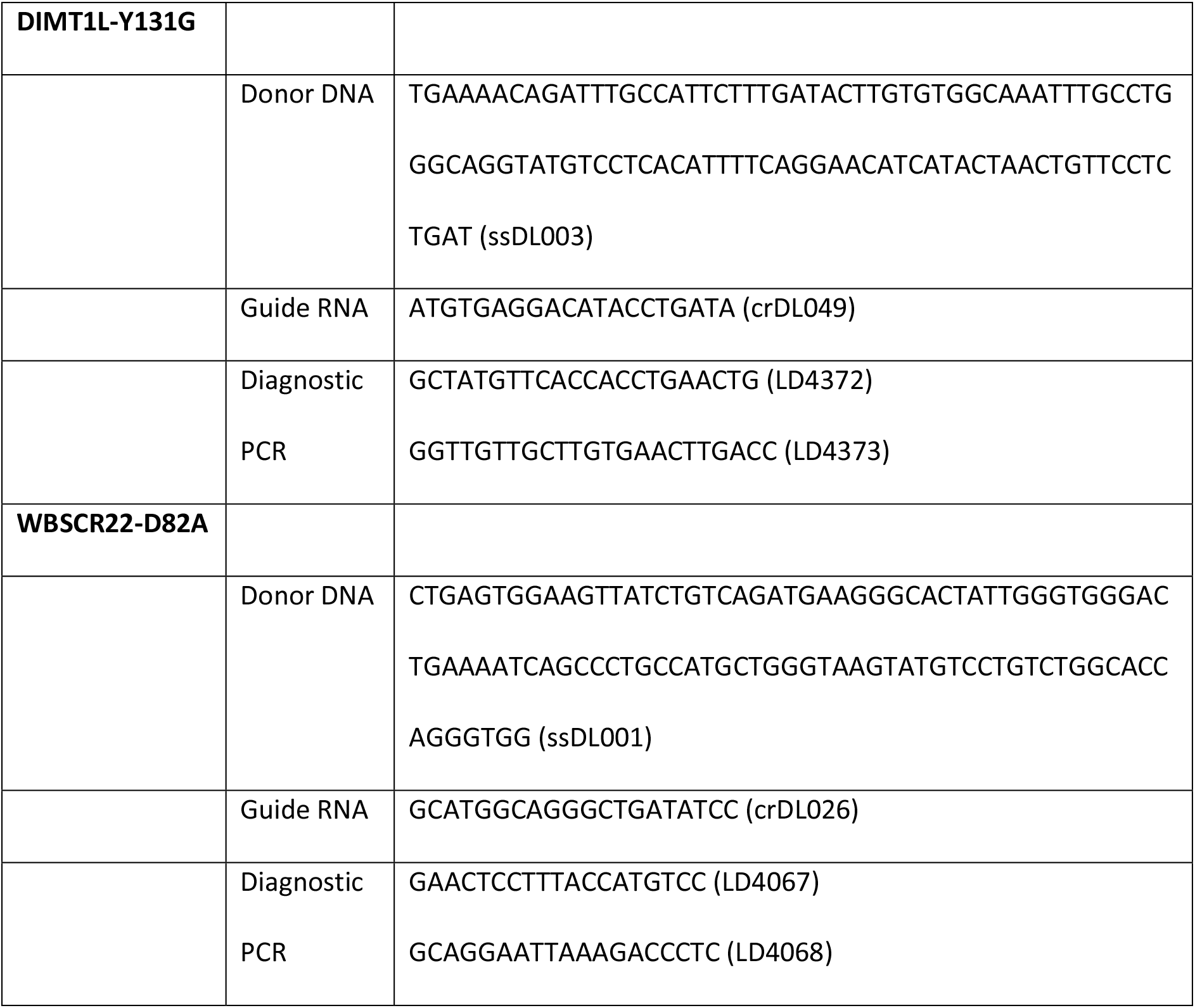
Sequence of oligonucleotides used for CRISPR-Cas9 knock-ins of methyltransferase mutations.

## Legends to Supplementary Figures

**Supplementary Figure 1:**
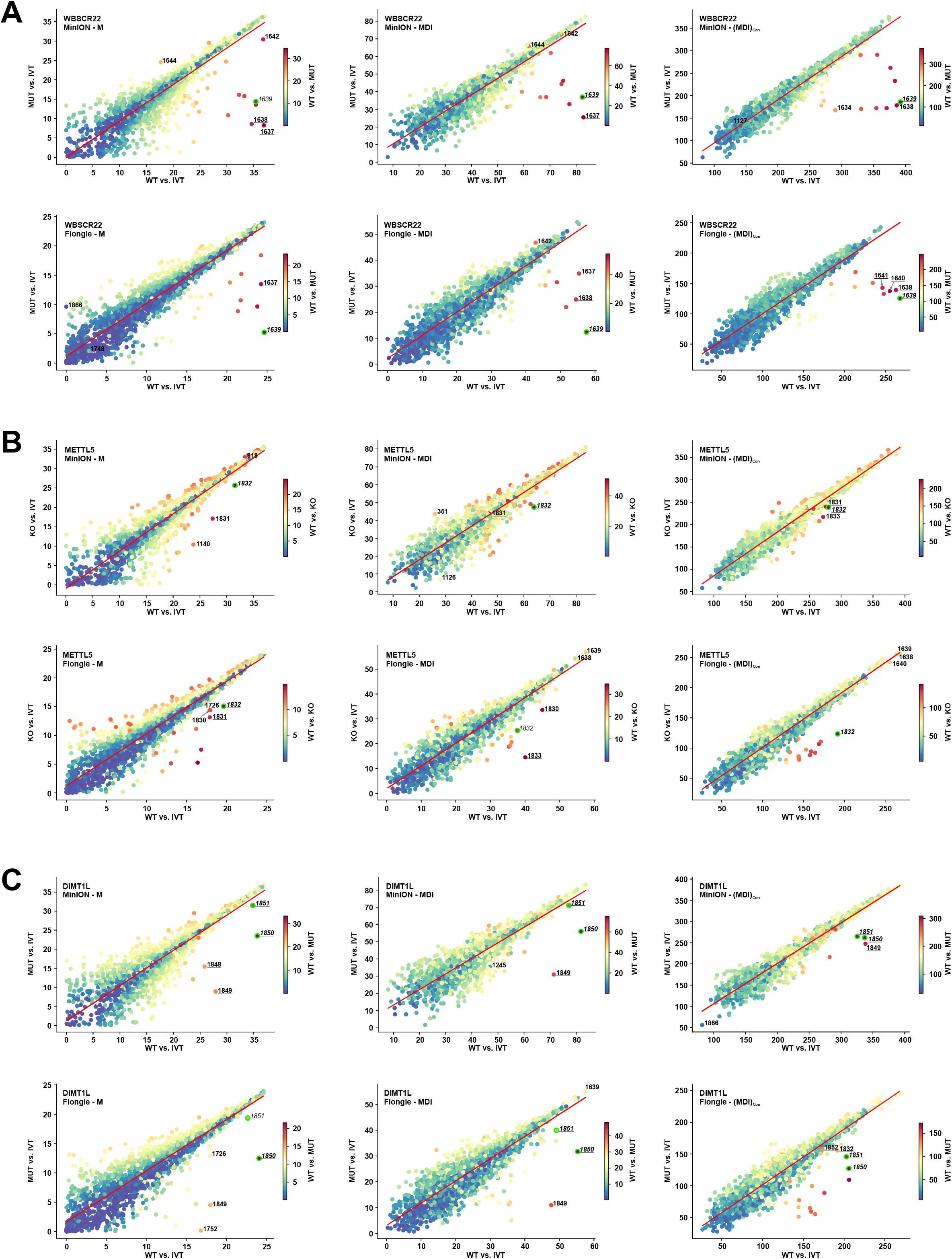
A) Coverage of Nanopore direct rRNA-seq of 18S rRNA from HCT116 WT cells sequenced on MinION R9.4.1 (top) or Flongle (bottom) flow cells. B) Coverage of Nanopore direct rRNA-seq of 28S rRNA from HCT116 WT cells sequenced on MinION R9.4.1 (top) or Flongle (bottom) flow cells.

**Supplementary Figure 2:**
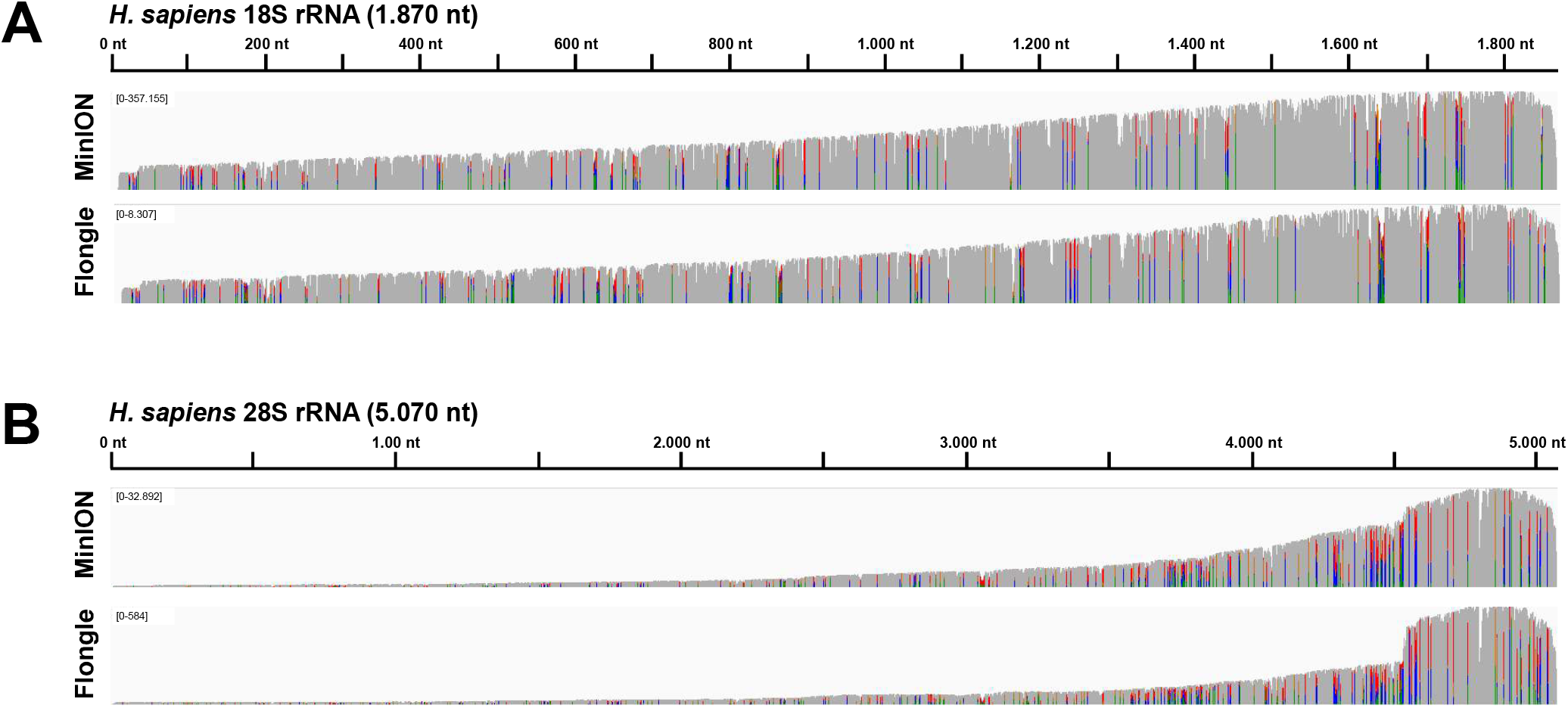
Analysis of genetic models by Nanopore direct rRNA-seq and JACUSA2. Shown are all pairwise comparisons of JACUSA2 call-2 from IVT, HCT116 WT and HCT116 KO/MUT for the Feature sets Mismatch (M, left panel), Mismatch+Deletion+Insertion (MDI, middle panel) and Mismatch+Deletion+Insertion for 5-mer context ((MDI)_Con_, right panel). The target site is indicated by a lime border, labeled are the target site (italics) and significant outliers detected by LOF (contamination value = 0.002, bold). Outliers in the 5-mer context are underlined. Analysis of MinION derived data is shown on the top of each panel, Flongle derived data are shown on the bottom. A) Analysis of 18S m^7^G_1639_ catalyzed by WBSCR22. B) Analysis of 18S m^6^A_1832_ catalyzed by METTL5. C) Analysis of 18S m^6^_2_A_1850_ m^6^_2_A_1851_ catalyzed by DIMT1L.

**Supplementary Figure 3:**
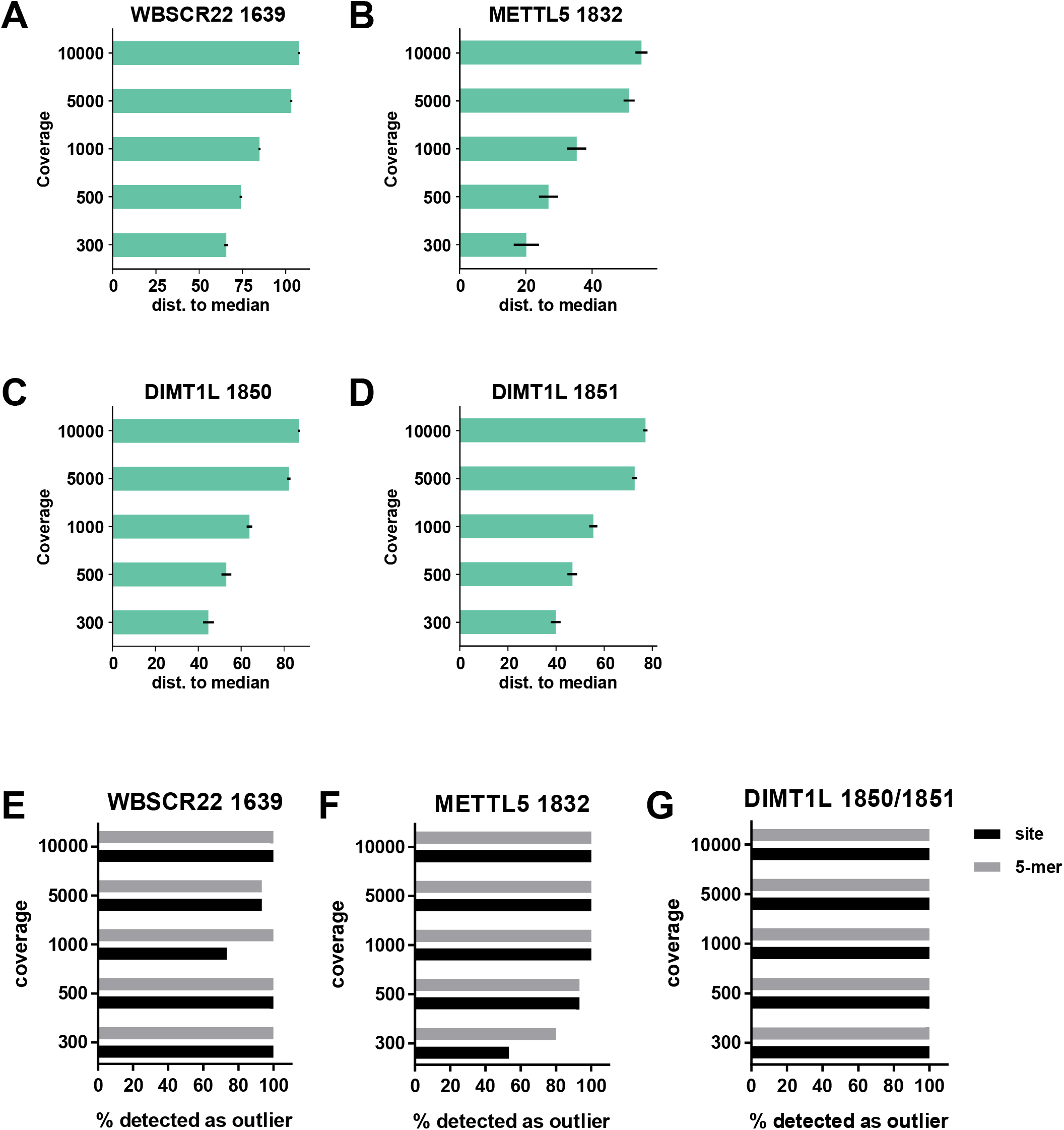
Detection of rRNA modifications with different read numbers. A) Distance of the JACUSA2 score for WBSCR22 m^7^G_1639_ to the median JACUSA2 score for the WT versus IVT comparison. Shown are the mean and the standard deviation from down sampling employing different seeds (n = 15). B) Analysis as in A for METTL5 m^6^A_1832_. C) Analysis as in A for DIMT1L m^6^_2_A_1850_. D) Analysis as in A for DIMT1L m^6^_2_A_1851_. E) Detection of the target site or the 5-mer context (target site in position 3) as LOF outlier (contamination value = 0.002) in down sampling with different read numbers for WBSCR22. F) Percentage of target sites/ 5-mer context detected as outlier as in E for METTL5. G) Percentage of target sites/ 5-mer context detected as outlier as in E for DIMT1L.

**Supplementary Figure 4:**
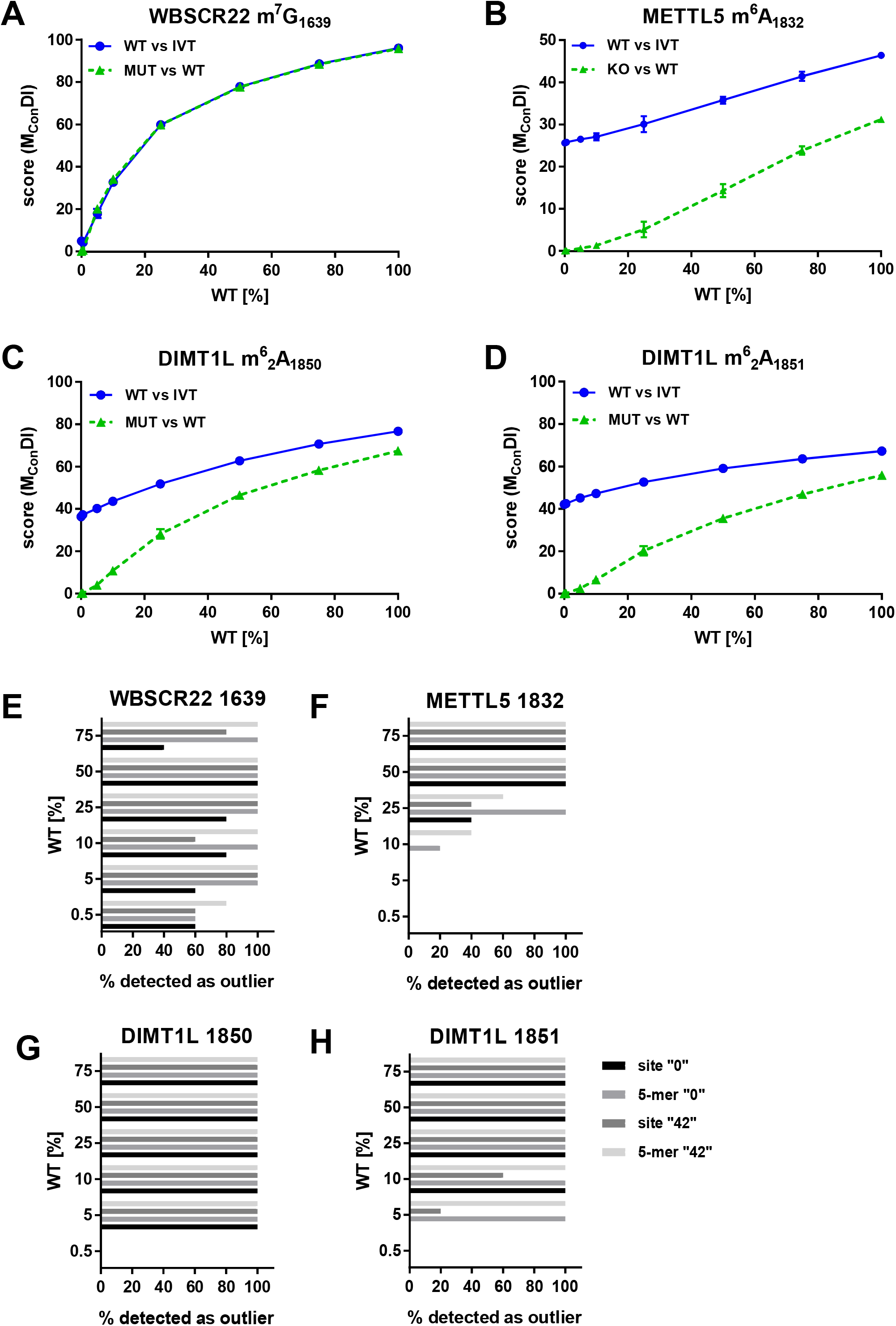
Analysis of the influence of modification level on the JACUS2A score. Analysis of the influence of modification level on the JACUS2A score. 1000 reads were downsampled with seed “42” from the MinION sequencing data shown in Figure 2. The “WT” sample was composed of modified (WT) and unmodified (KO/MUT) reads as indicated that were derived from the downsampled data with 5 different seeds. JACUSA2 call-2 analysis considering the M_Con_DI Feature set. The modification rate vs. the JACUSA2 score of the respective target site is shown (n =5). A) Analysis of m^7^G_1639_. B) Analysis of m^6^A_1832_. C) Analysis of m^6^_2_A_1850_. D) Analysis of m^6^_2_A_1851_. E) Detection of the target site or the 5-mer context (target site in position 3) as LOF outlier (contamination value = 0.002) in mixing analysis (n =5) from 1000 reads (down sampling with seed “0” or “42”) for WBSCR22. F) Outlier detection for METTL5 as in E. G) Outlier detection for DIMT1L 1850 as in E. H) Outlier detection for DIMT1L 1851 as in E.

**Supplementary Figure 5:**
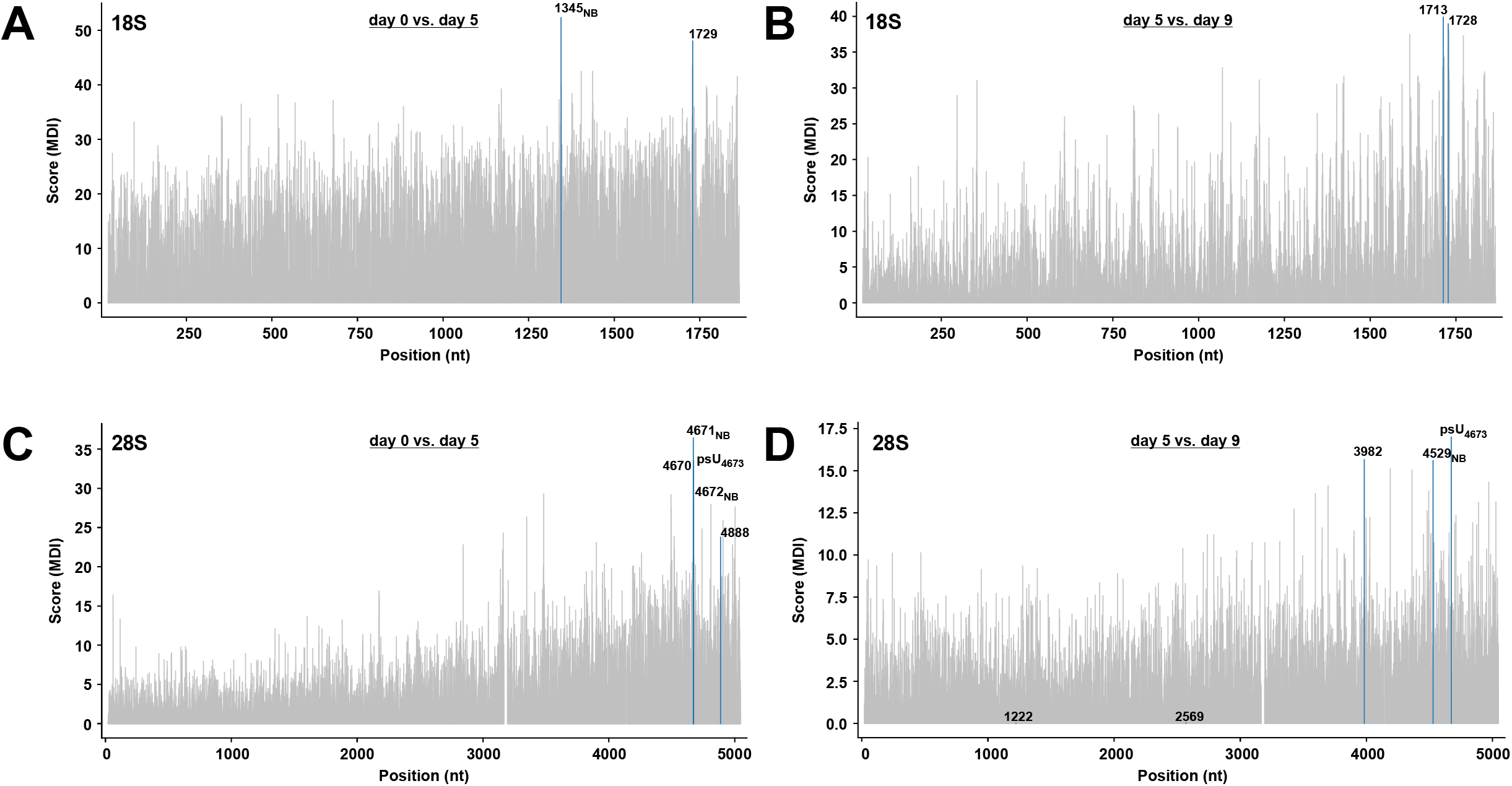
Analysis of rRNA modifications changes in cardiomyocyte differentiation. A) Bar plot of the pairwise comparison day 0 vs. day 5 for 18S rRNA. Significant outliers detected by LOF (contamination value 0.001) are labeled. B) Bar plot of the pairwise comparison day 5 vs. day 9 for 18S rRNA. Significant outliers detected by LOF (contamination value 0.001) are labeled. C) Bar plot of the pairwise comparison day 0 vs. day 5 for 28S rRNA. Significant outliers detected by LOF (contamination value 0.001) are labeled. D) Bar plot of the pairwise comparison day 5 vs. day 9 for 28S rRNA. Significant outliers detected by LOF (contamination value 0.001) are labeled.

